# Connecting Histopathology Imaging and Proteomics in Kidney Cancer through Machine Learning

**DOI:** 10.1101/756288

**Authors:** Francisco Azuaje, Sang-Yoon Kim, Daniel Perez Hernandez, Gunnar Dittmar

## Abstract

Proteomics data encode molecular features of diagnostic value and accurately reflect key underlying biological mechanisms in cancers. Histopathology imaging is a well-established clinical approach to cancer diagnosis. The predictive relationship between large-scale proteomics and H&E-stained histopathology images remains largely uncharacterized. Here we investigate such associations through the application of machine learning, including deep neural networks, to proteomics and histology imaging datasets generated by the Clinical Proteomic Tumor Analysis Consortium (CPTAC) from clear cell renal cell carcinoma patients. We report robust correlations between a set of diagnostic proteins and predictions generated by an imaging-based classification model. Proteins significantly correlated with the histology-based predictions are significantly implicated in immune responses, extracellular matrix reorganization and metabolism. Moreover, we showed that the genes encoding these proteins also reliably recapitulate the biological associations with imaging-derived predictions based on strong gene-protein expression correlations. Our findings offer novel insights into the integrative modeling of histology and omics data through machine learning, as well as the methodological basis for new research opportunities in this and other cancer types.

## 1. Introduction

Kidney cancer is one of the most common cancers worldwide accounting yearly for hundreds of thousands of deaths [1]. Clear cell renal cell carcinomas (CCRCC) is the most common subtype of kidney cancer representing ~75% of cases [2, 3]. Its diagnosis is typically incidental, e.g., as part of medical imaging tests unrelated to kidney problems, and ~30 % of patients with CCRCC eventually develop metastases even after removal of the kidney and other treatments [2]. Therefore, there is a need for developing new approaches to the understanding and early diagnosis of CCRCC.

Histopathology is a well-established technique for confirming diagnosis and subsequent sub-classification of kidney and other cancer types [4, 5]. Histopathology consists of the visual analysis of microscopic slides obtained from tissue samples typically stained with H&E (hematoxylin and eosin stains). This allows the pathologist to identify cellular patterns associated with the presence of cancer, its staging and potential clinical outcomes. Even when performed by well-trained experts, this task is time-consuming and not-always highly reproducible among pathologists [6, 7]. Moreover, in kidney and other cancers, the use of histological analysis for diagnostic purposes is often challenging because different cancer subtypes may share non-specific morphological patterns [2, 8]. Therefore, the accurate and robust analysis of large amounts of digitized histological slides for cancer diagnosis remains a key challenge in cancer research and clinical practice.

To address such challenges, different computational techniques have been proposed for analyzing histology images for diagnostic purposes in multiple cancers [9]. Such analyses have traditionally relied on the application of classification models, which process “handcrafted” (explicitly defined) image-derived features such as cell size, shape and pixel intensity distributions observed in full slides or selected slide patches [8, 9].

With the wider adoption of whole-slide high-content imaging and the increase in the volume of histology datasets, new opportunities have risen for the application of deep learning (DL) techniques [10]. Unlike previous generations of machine learning approaches, DL models based on convolutional neural networks (CNNs) can process raw intensity images and learn to automatically extract predictive features [11, 12]. The accuracy and potential clinical relevance of DL models for analyzing histology images for diagnostic and prognostic purposes have already been shown in different cancer research domains [13–15]. Thus, DL is expected to play a key role in the era of digital pathology and precision medicine [6, 16].

The analysis of large amounts of omics data, including transcriptomics and proteomics, has significantly advanced the molecular characterization of dozens of cancer types and offers deeper insights into their diagnosis, prognosis and treatment response assessment [17, 18]. This has been possible in large part because of consortia such as The Cancer Genome Atlas (TCGA) and the Clinical Proteomic Tumor Analysis Consortium (CPTAC) [18, 19]. For example, The TCGA recently reported a comprehensive analysis of multiple omics features of renal cancer and their associations with cancer subtypes and patient prognosis [3]. The study found that CCRCC tumors show elevated immune cell-specific gene expression in comparison to other kidney cancer sub-types.

The integration of omics and histopathology data has the potential to improve our understanding of the biological mechanisms underlying tumors, their detection and treatment [20]. Previous efforts to achieve these goals include the integration of H&E-stained tissue sections and genomic markers from patients diagnosed with gliomas [21]. CNNs were applied to analyze the images and predict patient survival, and the combination of such models with genomic biomarkers outperformed the current clinical prognosis approach [21]. In lung adenocarcinomas (LUAD), histopathology-derived features have been shown to correlate with omics-based classification, using gene and protein expression, and to improve patient survival prediction [22]. More recently, using LUAD and liver cancer datasets, the combination of gene expression and imaging features was also shown to improve patient prognosis [23]. In both investigations the histology-based prediction models processed inputs that represented handcrafted image-derived features reflecting specific cellular and sub-cellular morphological patterns. The application of DL models has also been demonstrated with TCGA-derived histopathology images and omics data. For instance, a CNN applied to whole-slide images showed a diagnostic performance comparable to that of pathologists, and was also capable of predicting the mutation status of commonly mutated genes in LUAD [24]. In breast cancer and using histopathology images, CNN-based models assigned patients to diagnostic attributes, e.g., tumor stage, and outperformed models based on transcriptomic data only [25].

Despite the progress achieved to date, such investigations tend to emphasize the implementation of histology-based models for improving classification accuracy. Moreover, the integrated analysis of such models with large-scale proteomics data have received relatively less attention in comparison to genomics and transcriptomics data. Deeper investigations of the association of histology imaging models and large-scale proteomics will not only improve our understanding of the predictive complementarity of such data sources, but also may offer the basis for more precise diagnostic systems. Here we address these research needs through the application of machine learning techniques, including DL models, for proteomics and imaging data. Based on the identification of correlations between image-based models and proteomics profiles, we generate hypotheses about the roles of proteins and biological processes in CCRCC, whose molecular activity can be accurately captured by histopathology imaging. Furthermore, to the best of our knowledge, we are the first team to systematically investigate the association of histology imaging and proteomics data in CCRCC using DL.

## 2. Methods

An overview of our research strategy is summarized in Figure 1A. Here we address the question of finding associations between diagnostic imaging and proteomics data. To achieve it, we analyzed histology images and proteomics data from hundreds of tumors and control samples. Machine learning models for distinguishing tumors from normal samples were built for each dataset independently (Figure 1B). Based on the resulting models, we investigated correlations between the diagnostic proteins and the image-based predictions. Using different databases containing annotations of biological processes and pathways, we detected statistically significant correlations that are relevant to cancer in general, and CCRCC in particular. Moreover, we investigated associations between mRNA obtained from the same patient cohort and the histology-based predictions, as well as between mRNA and their corresponding proteins.

**Figure 1.**
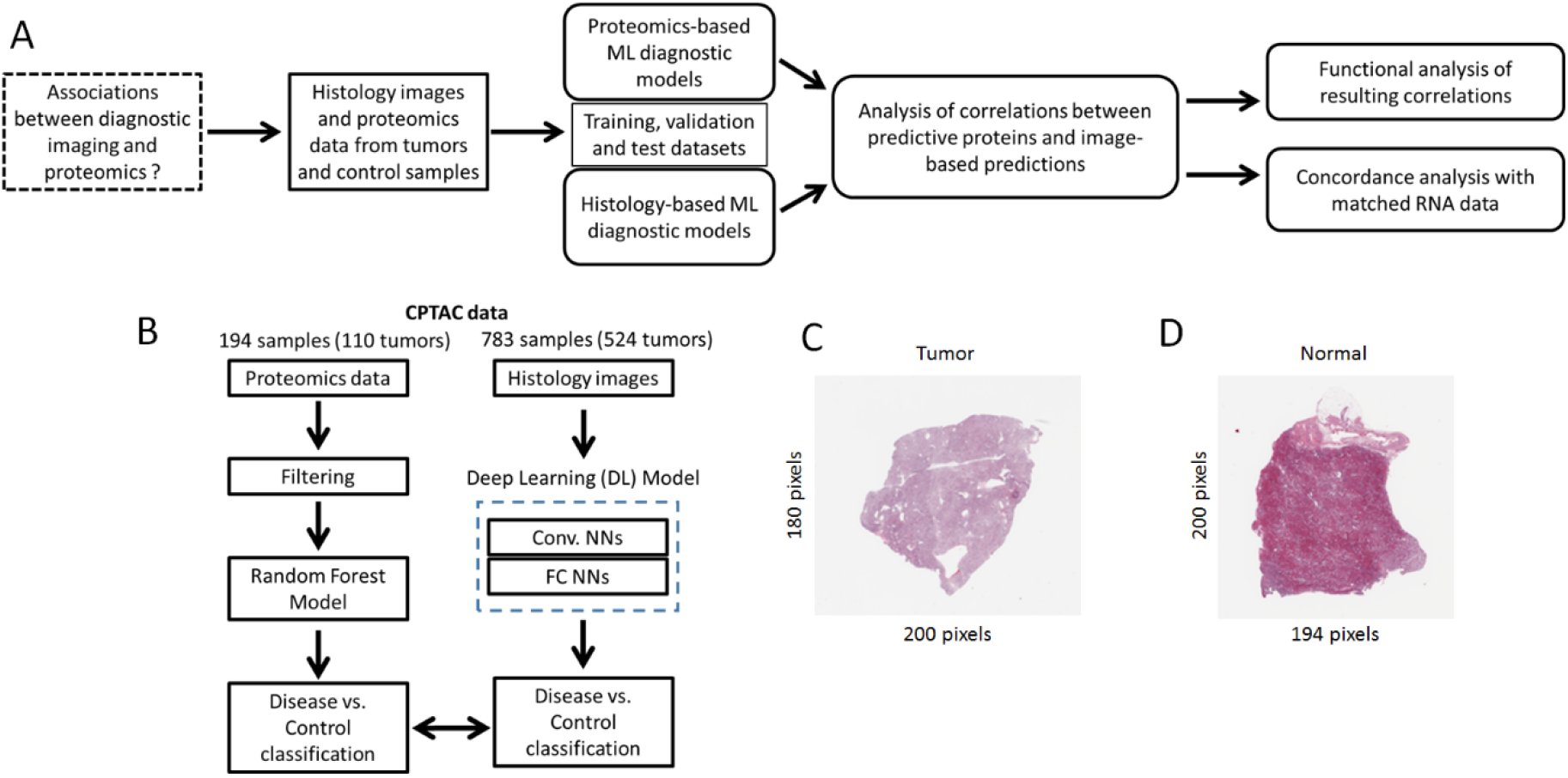
Overview of research strategy. A. Analytical and predictive modeling workflow implemented in this study. B. Focus on the implementation of diagnostic models based on proteomics and histology imaging. C., D. Examples of histology images obtained from tumor and normal samples respectively.

### 2.1. Datasets

The proteomics and histology datasets were generated by the CPTAC Clear Cell Renal Cell Carcinoma (CCRCC) Discovery Study [26]. The proteomics data, consisting of Tandem Mass Tags-10 (TMT10) experiments of 216 samples, were downloaded from the CPTAC Data Portal. This dataset included complete information for 9964 proteins measured in 194 samples (84 normal, 110 tumor samples), which are the focus of our investigation. The histology dataset was obtained from The Cancer Imaging Archive (TCIA), and included a total of 783 slide images (259 normal, 524 tumor, examples shown in Figures 1C and D). For some of the patients in this cohort, matching histology slides and proteomic samples (from the same patient) are available for investigating associations between proteomic- and image-based diagnostic models (details below). Before implementing diagnostic models, the proteomics dataset was pre-processed by selecting the LogRatio protein abundant column, and null values were replaced with zero. Raw histology images were fed into the DL models, and further processing at the pixel level was carried out during the model training process, as delineated next.

### 2.2. Diagnostic models

The proteomics-based diagnostic model was generated with a Random Forest (RF) classifier with default parameters and ntree = 500. As inputs to this model, we focused on the top-10% most variable proteins (based on their SD, i.e., 997 proteins) across all available samples. The RF model was trained, tested and its performance assessed with a 10-fold cross-validation (10-fold CV) sampling strategy. For both proteomics- and imaging-based models, we assessed their diagnostic performance using standard quality classification indicators: accuracy, precision, recall (sensitivity), F1 and AUC values.

The imaging-based diagnostic system consisted of a deep neural network architecture that combined: a CNN (the VGG16-CNN [27]), a regularized fully connected (FC) neural network and an output layer (OL). Because of the relatively small number of images (compared to typical large-scale datasets used in DL) and to reduce the computing times needed to train and test the models, we used a VGG16-CNN that was previously trained on more than 14 million generic images corresponding to 1000 image classes. Such a “transfer learning” is a well-established DL approach to extracting and re-using low-level image features across imaging application domains [10].

The histology imaging data were partitioned into training (181 normal and 366 tumor images), validation (52 normal and 105 tumor images) and test datasets (26 normal and 53 tumor images). These datasets were used for model generation, selection and independent evaluation respectively. To ensure an unbiased and robust analysis, we focused on the independent test dataset for implementing the proteomics-imaging integrative analysis. All the images were resized (to 224 x 224 pixels) and were input as 3-channel images to the DL model. To enable robust model building and reduce the risk of overfitting, images were randomly flipped and zoomed during training. The pre-trained VGG16-CNN was followed by a global average pooling layer, a fully connected network (128 units + ReLu activation) and a dropout layer to further minimize overfitting (rate = 0.2). Image classification was done with a 2-output (representing disease and control classes) using the softmax activation function to allow probabilistic classification. The FC and OL layers were optimized on the histology imaging data using the Adam optimization algorithm (lr = 0.001, decay = 0.0002), sparse categorical cross-entropy as loss function, with a maximum of 50 learning epochs and data batch size = 547.

### 2.3. Integrative data analysis

Correlations between protein expression and histology-based predictions (P-values generated by the DL diagnostic system) were calculated with the Pearson correlation coefficient. Out of the 79 images available in our independent test dataset, only 24 of them have patient-matched proteomics data. Functional enrichment analyses using GO, KEGG and Reactome annotations were implemented on the set of predictive proteins. To identify highly differentially enriched (Reactome) pathways in the proteomics data on the basis of their correlation with image-based predictions, we performed Gene Set Enrichment Analysis (GSEA) [28].

We also performed correlative, functional enrichment and GSEA analyses on mRNA data matched to the independent dataset, i.e., patients with proteomic, imaging and gene expression data. As the other datasets in this article, the gene expression data were generated by the CPTAC project (RNASeq) and analyses were applied to their FPKM expression values [29]. A total of 185 samples were available in the RNASeq dataset with matching proteomics data (including 110 tumors), and 9884 genes with corresponding proteins in the proteomics data. Among these data, 22 samples also have matched histology images.

### 2.4 Software and statistics

The proteomics-based RF classification model was implemented with the R packages caret and randomForest. The image-based DL classification model was implemented in Python using Pandas, NumPy, Matplotlib and Keras libraries. We applied one-sample t-tests for detecting statistical differences between matched data groups using R. The statistical significance of functional enrichment analysis and GSEA was estimated with Benjamini-Hochberg adjusted p-values. Additional data processing and visualization tasks were completed with R packages: fgsea, Rtsne, ggplot2 and complexHeatmap.

## 3. Results

### 3.1. A proteomics-based classification model accurately detects CCRCC

Before implementing the proteomics-based classifier, we investigated the sample discrimination potential of the top-variable 997 proteins using an unsupervised classification algorithm. We found that this set of proteins effectively segregates disease and normal samples into clearly separated clusters (t-SNE mapping, Figure S1). Interestingly, when using the full set of proteins available in the dataset, we obtained a relatively good segregation of samples as well: Only 3 normal samples were clustered closer to tumor samples than to normals (Figure S1). These observations corroborate both the quality and diagnostic potential of the proteomics dataset, in general, and of our selected set of 997 proteomic markers, in particular.

The proteomics-based RF classification model was capable of distinguishing between CCRCC and normal samples with an overall accuracy of 0.98 (10-fold CV results), as well as high sensitivities and specificities (0.97 and 0.99 respectively). This also resulted in high F1 and AUC values (0.98 and 0.99, 10-fold CV results), which offer further evidence of the powerful diagnostic capacity of our proteomics-based classification model.

### 3.2. A histology-based classification model accurately detects CCRCC

The histology-based prediction (DL) model was trained using the transfer learning and network adaptation strategy detailed in Methods. The training process was implemented to learn the parameters of the FC and OL layers of our DL model, while keeping the (transferred learning) parameters of the CNN frozen. The resulting models consistently reported classification accuracies between 0.98 and 0.99 (on the training dataset), and between 0.81 and 0.88 when evaluated on a separate validation dataset (Methods). Such classification performance was observed when training our DL model during 50 epochs. A relative high classification performance was also obtained on the validation dataset for fewer training epochs: Accuracies between 0.83 and 0.85 (for 3 and 20 training epochs respectively). To reduce the risk of model overfitting and decrease the time needed for training and evaluating models, we selected a DL model trained with 3 learning epochs and the parameters specified in Methods.

The selected model was then applied to the independent test dataset of histology images. Our histology-based classification model was capable of distinguishing between CCRCC and normal samples with an accuracy of 0.95 on the test dataset, as well as with high sensitivities and specificities (1 and 0.93 respectively). This also resulted in high F1 and AUC values (both equal to 0.92), which further indicates the solid diagnostic capacity of our model. The model actually only misclassified 4 images out of 79 test images: 4 normal images predicted as tumors.

### 3.3. Proteomic markers are correlated with histology-based predictions

The previous section’s findings motivated us to investigate in depth the relationship between the proteomic markers and the histology-based model predictions. Knowing that the proteomics data represent a strong source for accurately classifying normal vs. tumor samples, a key question is how such predictive features relate to the image-based predictions. To answer this question, first we calculated correlations between each protein in our test dataset of 24 samples (14 tumors, 10 normal samples) and their corresponding image-based predictions (P-values of assigning a sample to the tumor class). Also using hierarchical clustering, we further demonstrated that the protein expression data are sufficient to accurately separate tumor from normal samples (Figure 2A). Moreover, these proteins can be grouped in terms of their (expression) correlations with the image-based predictions (plot shown on left side of heatmap, Figure 2A). In particular, the histology-derived predictions are strongly associated, either highly positively-or anti-correlated, with a sub-set of protein markers (Figure 2B).

**Figure 2.**
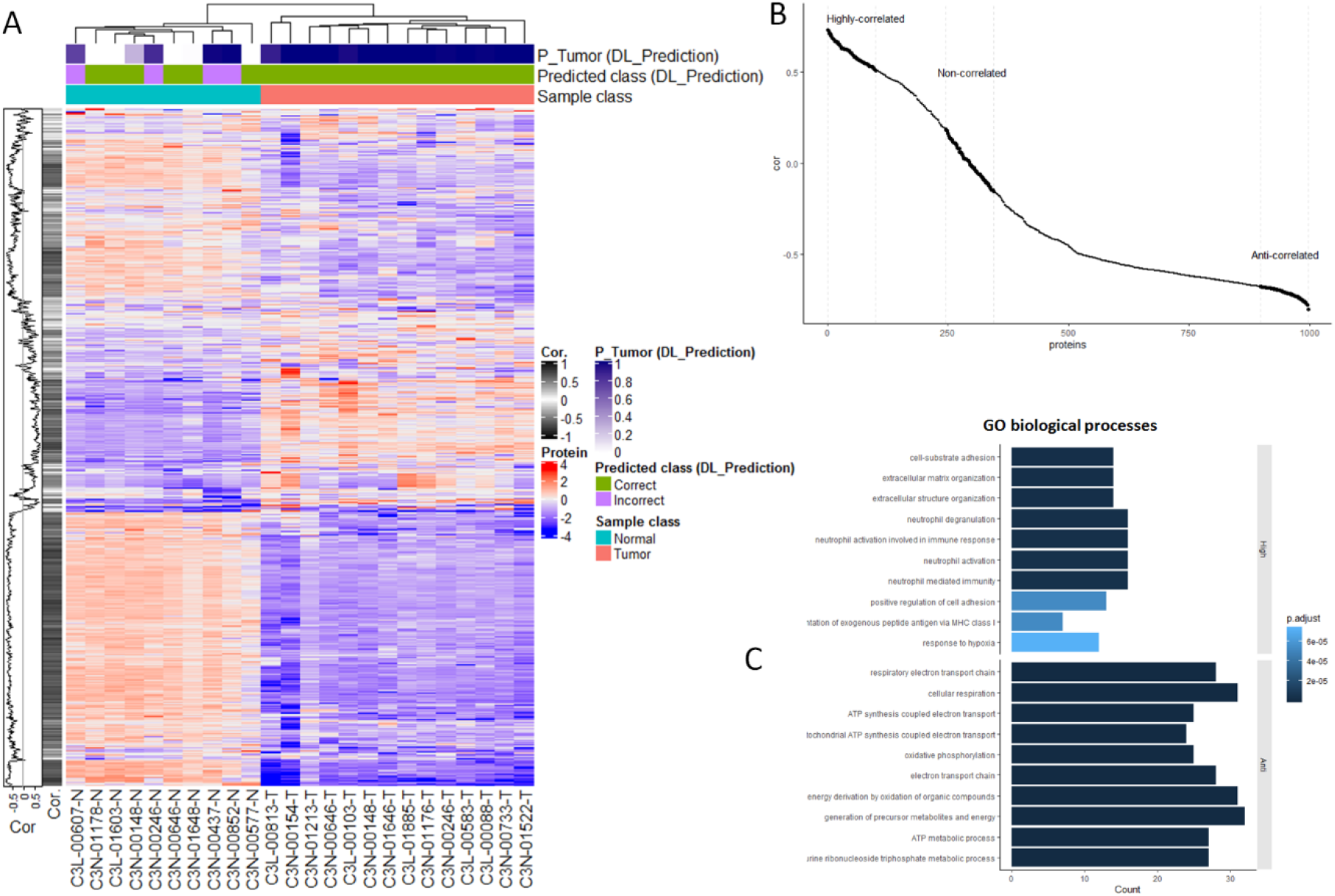
Proteomic markers are correlated with histology-based predictions. A. Heatmap of proteomics data (vertical axis: Expression values for 997 proteins) measured in 24 samples in our independent test set (horizontal axis). The top of the heatmap specifies the true and predicted classes (normal and tumor samples), as well as the corresponding P-values of assigning a sample to the tumor class, as predicted by our histology-based DL model. The plot on the left side of the heatmap depicts the correlations between each protein and the predictions generated by the histology-based DL model (P-values of tumor classification). B. Plot showing the correlations between each protein expression and the predictions generated by the histology-based DL model (P-values of assigning an image to the tumor class). Correlation values are ranked from the highest positive to lowest negative (anti-correlated) values. C. GO enrichment analysis of proteins highly positively- and anti-correlated with the histology-based predictions. Bars indicate the magnitude of the enrichment scores, and adjusted P-values of the enrichments are color coded. Statistically enriched GO terms were not detected for proteins whose expression values were weakly correlated with histology-based predictions (i.e., those with correlations around 0).

A closer examination of these relationships showed that the proteins that are either highly positively- or anti-correlated with histology-based predictions are significantly enriched in a diversity of biological processes (Figures 2C and S2, GO terms and KEGG pathways respectively). In the case of proteins that are highly positively correlated with the image-based predictions, such an enrichment includes processes relevant to cell adhesion, extracellular organization and immune responses (Figure 2C). Proteins that are strongly anti-correlated with image predictions are significantly associated with several respiratory and metabolic processes. Unlike highly positively and anti-correlated proteins, weakly correlated proteins, i.e., those with correlations around 0 (Figure 2B), are not statistically associated with specific biological processes.

### 3.4. Independent verification of biological associations

Using an independent database of annotated molecular pathways (Reactome) and an alternative enrichment analysis technique (GSEA), we found again that proteins either strongly positively- or anti-correlated with histology-based predictions are significantly enriched in a variety of cancer-relevant molecular pathways (Figure 3A). Unlike the analysis reported above, here we considered the actual levels of the observed correlations between the proteomic data and the histology-based predictions for detecting significant functional enrichments.

**Figure 3.**
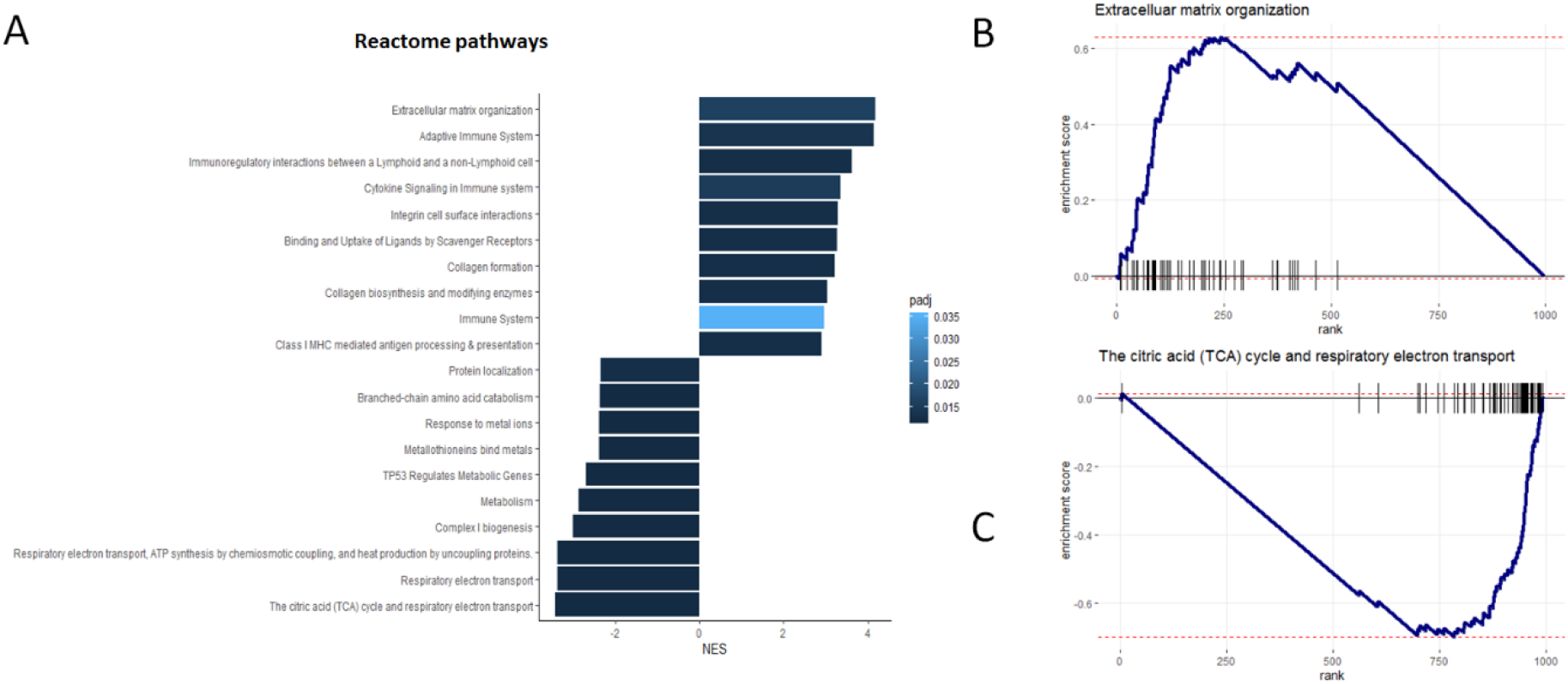
Proteins either strongly positively- or anti-correlated with histology-based predictions are significantly enriched in cancer-relevant molecular pathways. A. List of Reactome pathways that are significantly associated with the proteins on the basis of their correlations with histology-based predictions, as detected by GSEA (Methods). Bars indicate the magnitude of the enrichment scores, and adjusted P-values of the enrichments are color coded. B. and C. show examples of pathways significantly associated with proteins highly positively- and anti-correlated with histology-based predictions respectively. In B. and C. proteins are ranked according to their correlations with the histology-based predictions, from highest to lowest, and pathway enrichment scores were estimated with GSEA.

We verified that proteins that are positively correlated with the imaging-based predictions are also statistically associated with molecular pathways relevant to extracellular organization and immune responses (Figure 3A and 3B). Conversely, we found that proteins that are anti-correlated with histology-based predictions are significantly associated with respiratory and metabolic pathways (Figure 3A and 3C). These findings provide additional supporting evidence of the direct connection between proteomics markers and histology-based predictions, as well as of their biological meaning in the specific context of CCRCC.

### 3.5. Genes are highly correlated with proteomic markers and imaging-based predictions

Next, we analyzed the concordance between proteins and their coding RNAs on the basis of their expression values. This analysis was applied to a set of 22 samples (14 tumor and 8 normal samples) with matched proteomics, gene expression and imaging data available. Figure 4 displays a global view of the correlations between these datasets and the histology-based predictions independently. To facilitate a comparative visualization of major trends, in each plot the rows show proteins (Figure 4A) and their corresponding genes (Figure 4B) in full alignment. This analysis first indicates that, as the proteomics data, the gene expression data are sufficiently informative to perfectly separate tumors from normal samples (Figure 4). Moreover, as observed in the case of the proteomics data, genes can also be meaningfully ranked on the basis of their correlations with the histology-based predictions (see correlation plots on the left side of each heatmap, Figure 4). The latter includes RNAs highly positively- and anti-correlated with the histology-based predictions (Figures 4 and S3). Also, as in the case of the proteomics data, such genes are significantly enriched in biological processes (Figure S3): immune responses and extracellular organization (for genes highly positively correlated with histology-based predictions), and metabolic processes (for genes anti-correlated with histology-based predictions).

**Figure 4.**
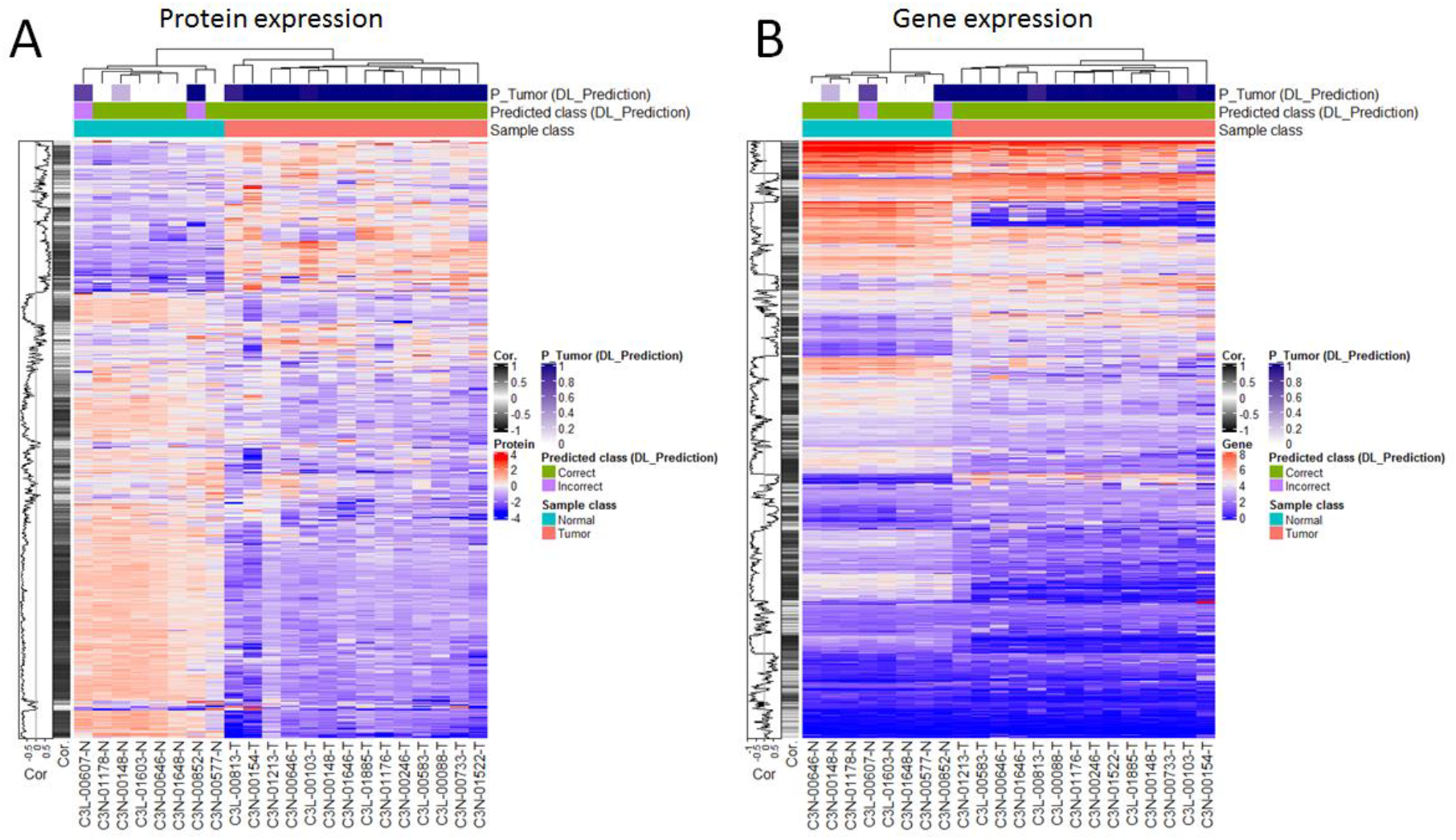
Analysis of the correlations between the proteomics data and their encoding genes, and between each dataset and the histology-based predictions. A. Focus on the proteomics data. B. Focus on the gene expression data. In each plot, the rows show proteins (A) and their corresponding coding genes (B) in alignment to facilitate comparative visualization, i.e., each row in each heatmap refers to a protein and its coding gene. Analysis performed on 22 samples with matched proteomics, gene expression and imaging data available.

A deeper analysis of these datasets (995 proteins with their corresponding gene expression data) showed strong correlations between protein and gene expression (median absolute Pearson correlation, *r* = 0.76). This correlation was statistically higher than that observed when all the proteins available in the dataset (N = 9984 proteins with corresponding gene expression data) are considered (*r* = 0.76 vs. 0.47, P < 2.2E-16, Figure 5A). Moreover, we found that the correlations between protein expression and image-based predictions are also concordant with the correlations between gene expression and image-based predictions, in particular for the strongest positive and negative correlations observed in each correlation setting (Figure 5B).

**Figure 5.**
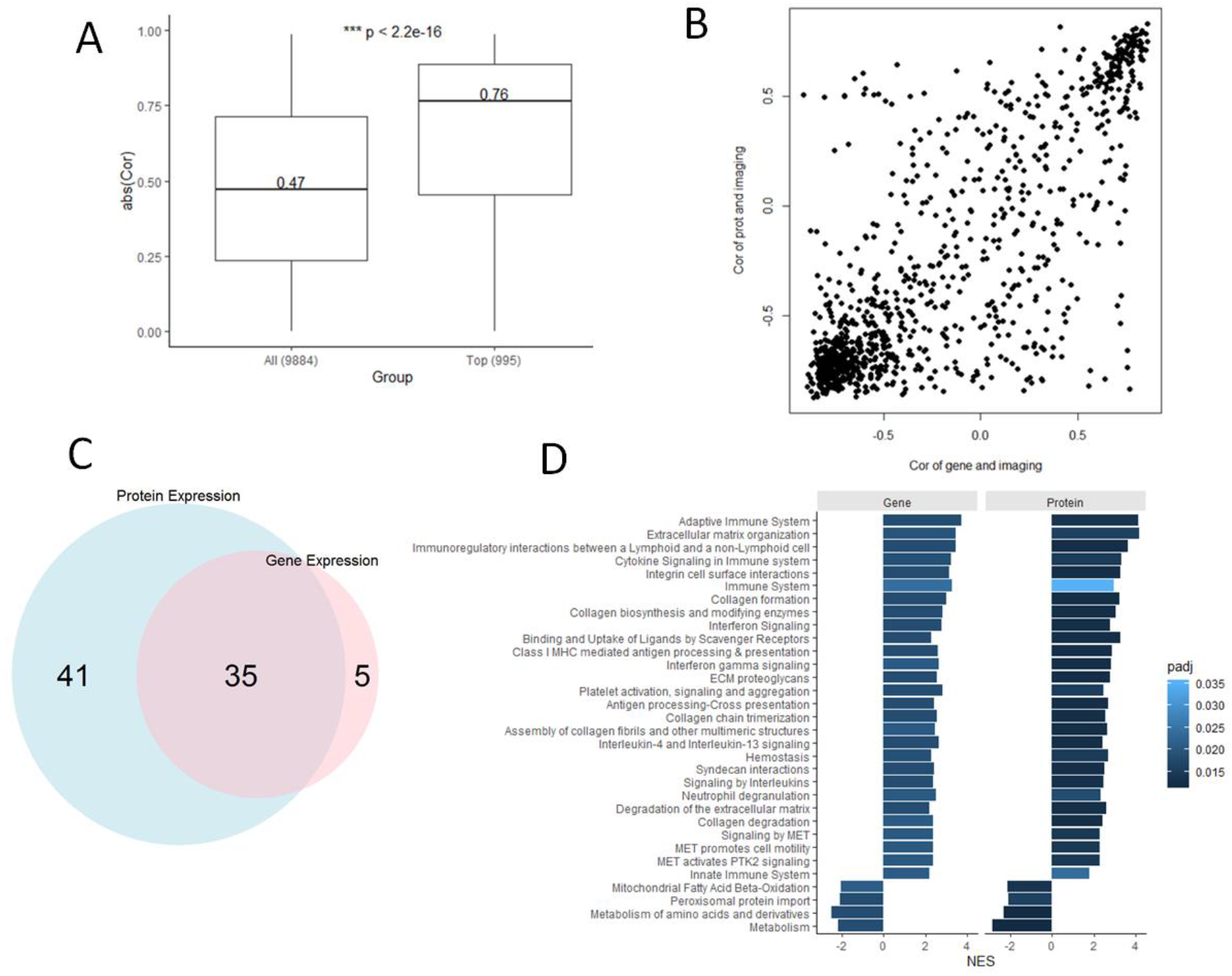
Protein and gene expression data are highly concordant in the diagnosis of CCRCC. A. Box plot of the gene-protein expression correlations observed in our set of 995 proteins, compared to distribution of correlation values observed in the full set of proteomics dataset with available gene expression data. B. Correlation plot of protein expression-image prediction correlations (vertical axis) vs. gene expression-image prediction correlations (horizontal axis). C. Number of overlapping molecular (Reactome) pathways statistically detected (with GSEA) in the protein and gene expression datasets independently, on the basis of their correlations with the image-based predictions. D. List of significantly enriched 35 pathways shared in common by the protein and gene expression datasets. Bars indicate the magnitude of the enrichment scores, and adjusted P-values of the enrichments are color coded.

GSEA of the proteins and genes separately, ranked by their correlations with the image-based predictions, resulted in 35 statistically enriched molecular pathways that were detected by both datasets independently (Figure 5C). This shared set of functional associations included 31 pathways relevant to different immune and extracellular matrix organization processes with positive enrichment scores, i.e., the correlations of protein (and gene expression) with image-based predictions are also positively correlated with the activity of these pathways (Figure 5D). Conversely, there are 4 pathways relevant to different metabolic processes with negative enrichment scores (Figure 5D). The latter means that image-based predictions that are not positively correlated with protein and gene expression are similarly anti-correlated with the activity of these 4 pathways.

To further assess the relevance of the correlations between protein (and gene) expression and image-based predictions, we investigated whether only the correlations between protein and gene expression would be sufficient to detect the above-identified molecular mechanisms independently of the image-derived prediction information. This analysis was done by ranking the 995 proteins on the basis of their expression correlations with their corresponding encoding genes, i.e., from the highest to the lowest protein-gene expression correlation pairs, followed by GSEA applied to the obtained ranking. This analysis did not result in any significant pathway enrichments for the set of 995 proteins, though as expected a variety of pathway enrichments were found when using the full set of 9884 proteins (Figure S4). These results confirm that histology imaging-based predictions can reliably capture information relevant to immune responses and metabolic processes, as encoded in both the proteomics and transcriptomics data.

## 4. Discussion

Our research addressed the problem of integrating histopathology- and proteomics-based diagnostic models through machine learning approaches. This challenge is important for systematically determining molecular features that can be accurately captured by pathology-based diagnostic models. Although our proteomic- and pathology-based models do not show perfect classification capacity, they are sufficiently accurate for investigating predictive relationships between them, as well as for establishing commonalities and complementarities at the functional level.

Using CCRCC as a novel study case, we elucidated the correlation of the diagnostic proteomics data with the predictions generated by the histology-based diagnostic model. This analysis demonstrated that, on the basis of their expression, a set of proteins are strongly correlated with the image-derived predictions. Using multiple annotation datasets and statistical analyses, we also showed how these correlations are significantly linked to specific biological processes relevant to the emergence and development of cancer. More specifically, we showed how our histology-based diagnostic model accurately captures predictive features in the proteomics dataset that are implicated in immune responses and extracellular matrix re-organization. These associations are also relevant in light of recent findings by the TCGA showing that CCRCC tumors are characterized by elevated immune activity [3]. Conversely, we showed how anti-correlations between proteomics and histology models are reflective of metabolic processes. Furthermore, we showed that gene expression data can also very closely recapitulate these biological associations based on their strong correlation with the proteomics data. These findings are useful not only for understanding novel ways to integrate these data types for predictive purposes, but also for generating hypotheses about the mechanisms underlying patient-specific classifications.

Although our study offers novel and relevant insights into the integration of histology and proteomics data through the application of machine learning, it shows some limitations that will merit future consideration. First, our study is limited by our focus on a single patient cohort of CCRCC patients. Additional validations on datasets obtained from independent cohorts may further demonstrate the clinical relevance of our diagnostic models and their integrative analysis, and will also enable wider investigations of variations related to different clinical factors, such as gender and tumor subtypes. Nevertheless, our study provides a solid basis for further investigations based on the analysis of carefully annotated datasets obtained from a CPTAC reference cohort. Our study is also limited by the relatively small amounts of data, in particular those needed for independently validating our models on matched histology and proteomics data from the same patients. Although the CPTAC currently offers the largest amount of data combining histology and proteomics data for CCRCC research, further validations with cohorts of different sizes are needed.

To conclude, our study presented a systematic investigation of the association of histopathology and proteomics data in a diagnostic setting. The resulting models and insights are relevant for understanding the predictive interplay between these datasets, as well as their informational complementarities at the molecular level. Furthermore, the proposed integrative analysis approach is applicable to other investigations with different tumors or omic data types.

## Author Contributions

conceptualization, F.A.; methodology, F.A., D.P.H, G.D.; software, F.A., S.Y.K.; investigation, F.A., S.Y.K., D.P.H, G.D.; writing—original draft preparation, F.A.; writing—review and editing, F.A., S.Y.K., D.P.H, G.D.

## Acknowledgments

Data used in this publication were generated by the National Cancer Institute Clinical Proteomic Tumor Analysis Consortium (CPTAC). We thank the developers of the Keras deep learning library and the VGG16-CNN for making their APIs open and freely available.

---

**Figure S1.**
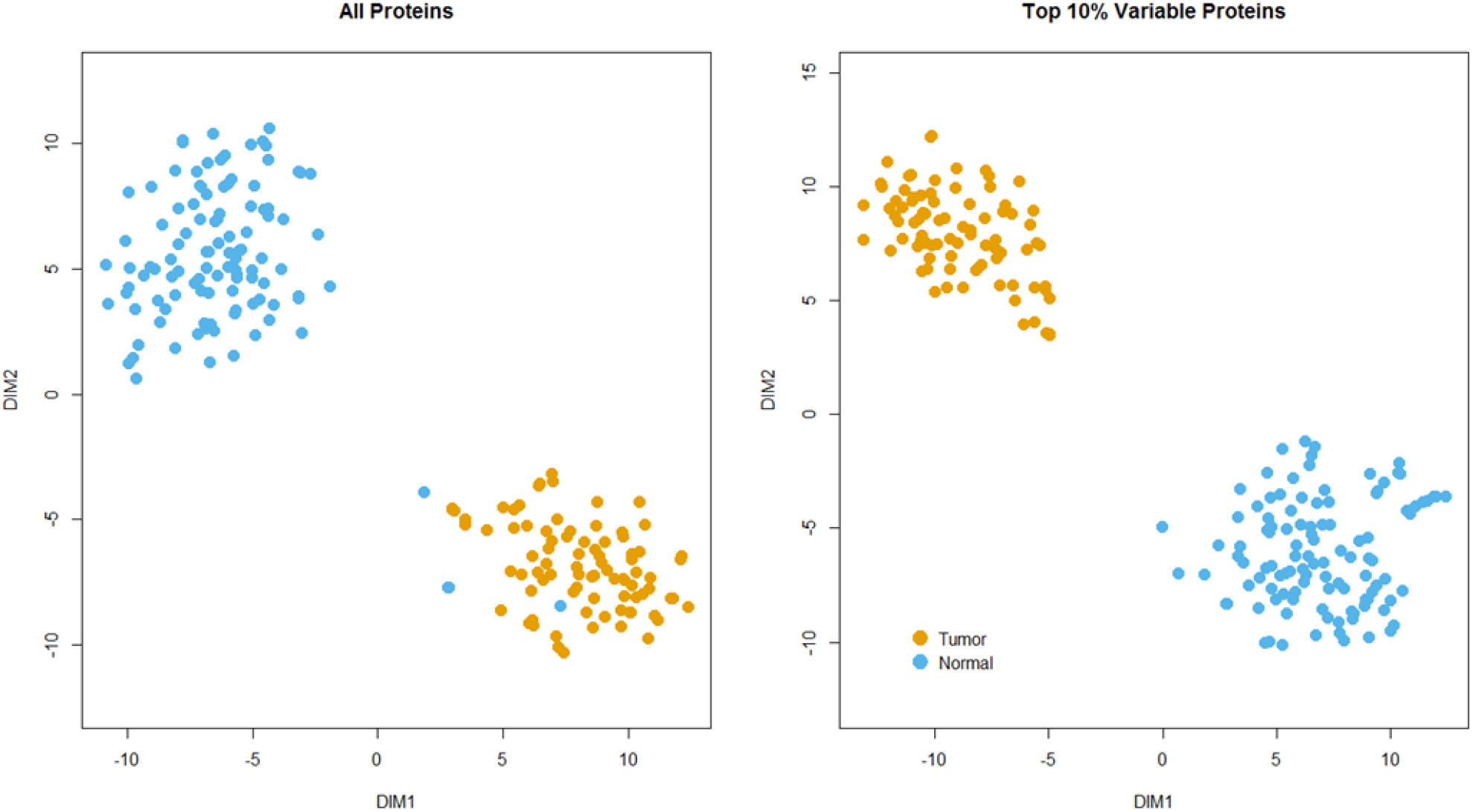
Unsupervised exploration of the sample discrimination potential of the CPTAC-CCRCC proteomics dataset. Both plots show 2D maps of the projected transformations generated by the t-SNE algorithm. Left plot was obtained when using the full set of proteins available in the CCRCC dataset (9964 proteins). The right plot was obtained when using the top 10% most variable proteins in the dataset.

**Figure S2.**
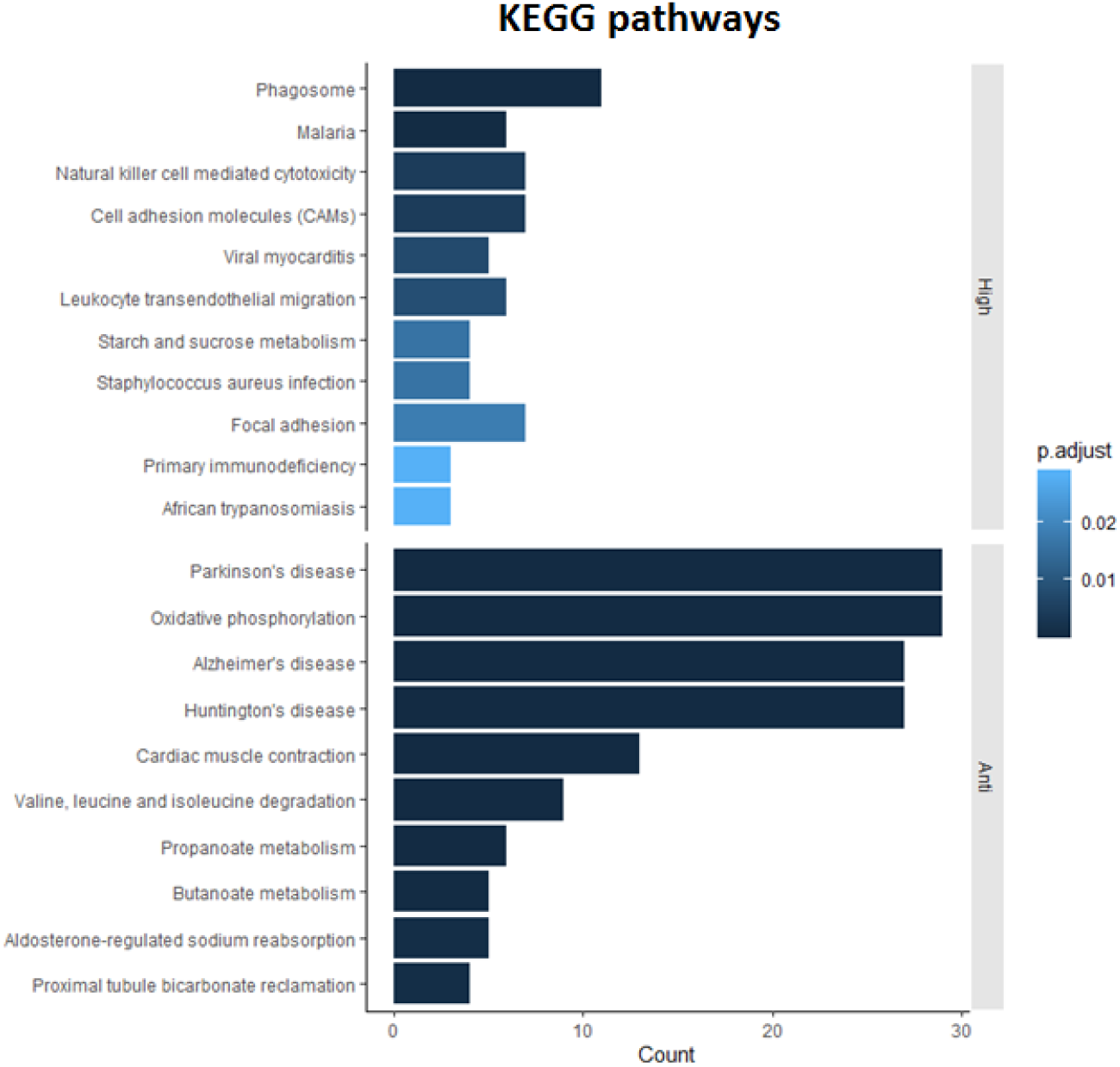
KEGG pathway enrichments of the proteins that are either highly positively-or anti-correlated with histology-based predictions. Bars indicate the magnitude of the enrichment scores, and adjusted P-values of the enrichments are color coded.

**Figure S3.**
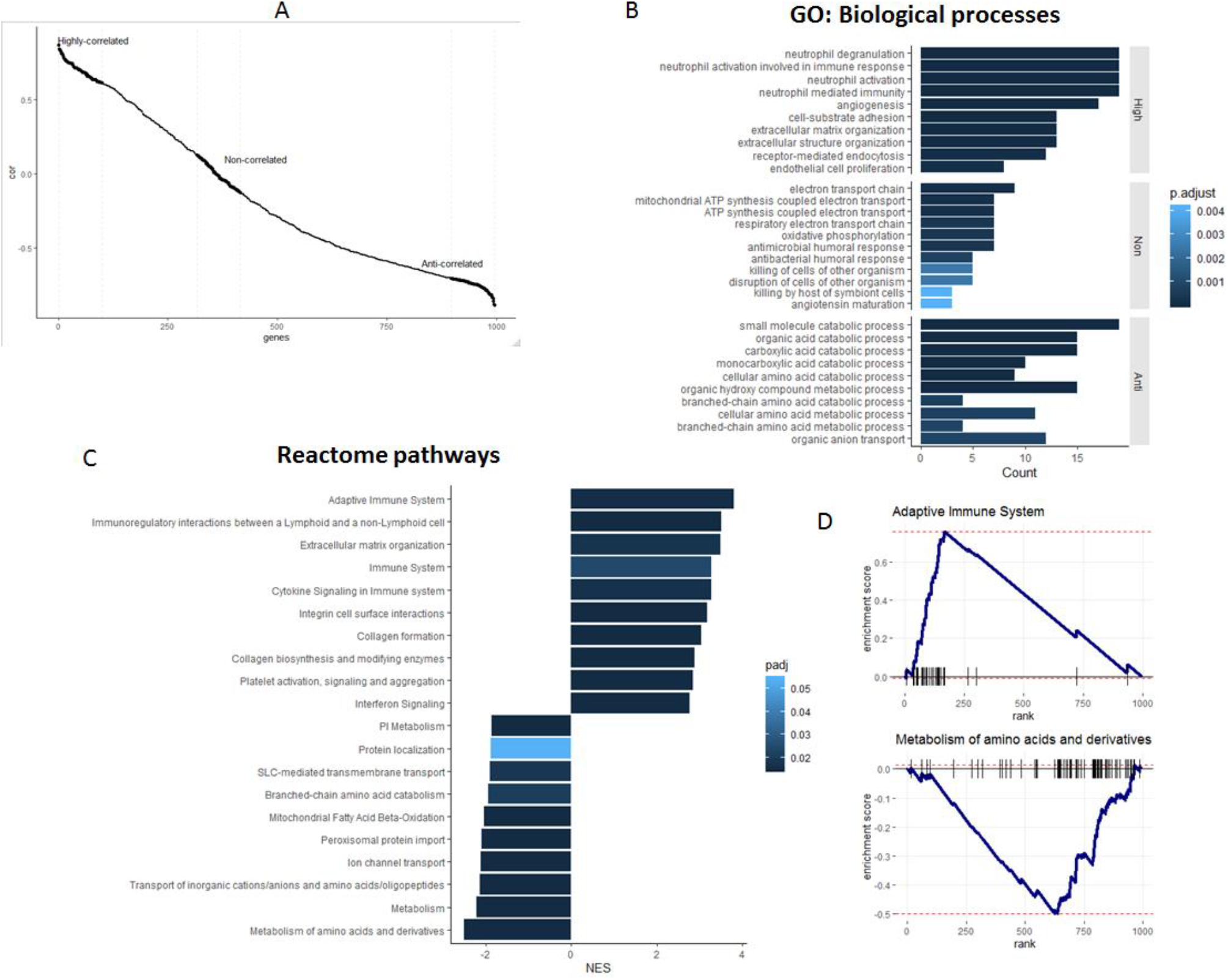
Analysis of correlations between RNAs (encoding the diagnostic set of proteins) and the histology-based model predictions. A. Correlation plot of gene expression values and image-based predictions (P-values of assigning an image to the tumor class). B. GO enrichment analysis of proteins highly positively-, anti- and not correlated with histology-based predictions. C. List of Reactome pathways that are significantly associated with the genes on the basis of their correlations with histology-based predictions, as detected by GSEA. D. Examples of pathways significantly associated with proteins highly positively- and anti-correlated with histology-based predictions respectively. In B and C: Bars indicate the magnitude of the enrichment scores, and adjusted P-values of the enrichments are color coded.

**Figure S4.**
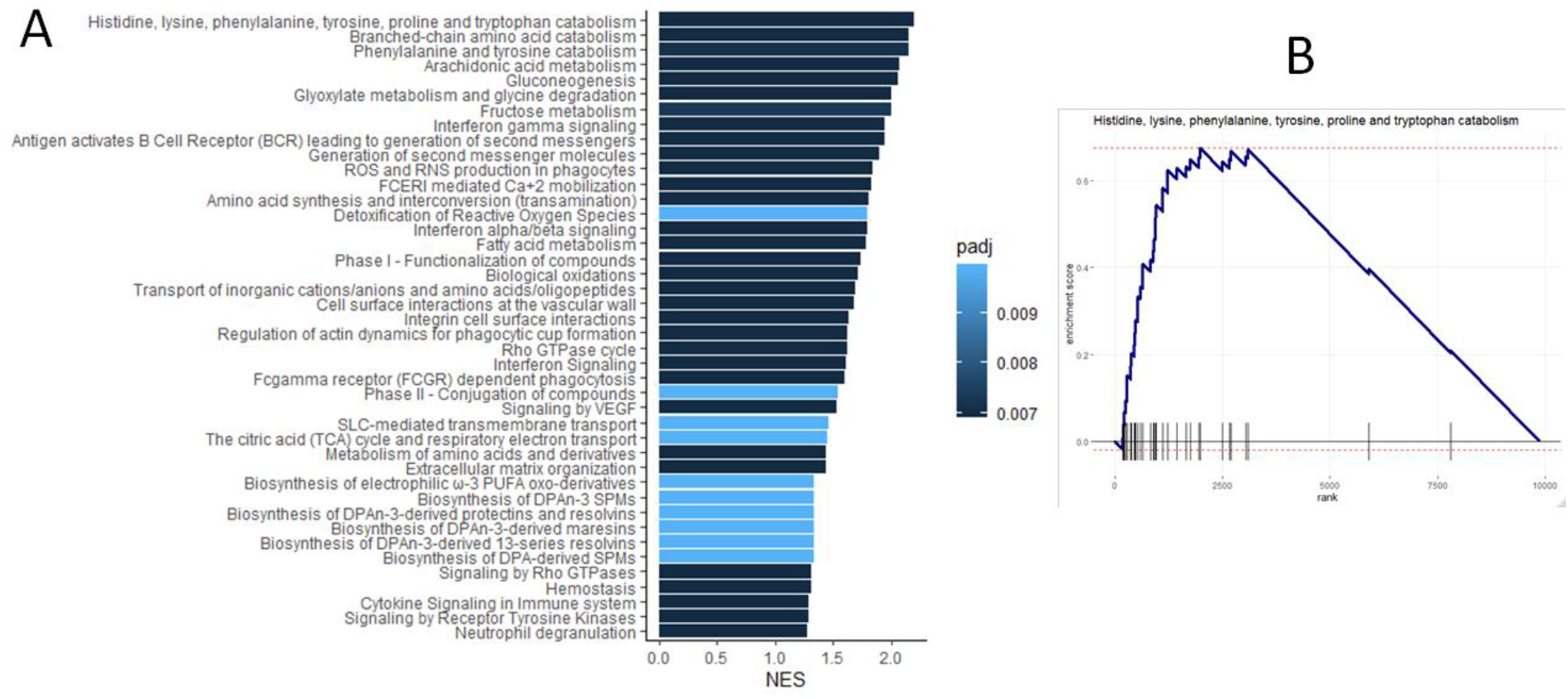
Reactome pathways enriched in the full set of 9884 genes available in the CCRCC (RNASeq) dataset. A. List of statistically detected enrichments identified by GSEA. B. Example of a significantly enriched molecular pathway. Genes were ranked on the basis of their expression correlations with their protein expression values, and GSEA estimated pathway enrichment scores. Using the same analysis, no significant enrichments were obtained when focusing on the set of 995 diagnostic proteins.

## References

1. WHO. Cancer Today. Available from: (https://gco.iarc.fr/today).

2. Hsieh, J.J., et al., Renal cell carcinoma. Nat Rev Dis Primers, 2017. 3: p. 17009.

3. Linehan, W.M. and C.J. Ricketts, The Cancer Genome Atlas of renal cell carcinoma: findings and clinical implications. Nat Rev Urol, 2019.

4. Turk, J.L., Rudolf Virchow--father of cellular pathology. J R Soc Med, 1993. 86(12): p. 688–9.

5. Fischer, A.H., et al., Hematoxylin and eosin staining of tissue and cell sections. CSH Protoc, 2008. 2008: p. pdb.prot4986.

6. Djuric, U., et al., Precision histology: how deep learning is poised to revitalize histomorphology for personalized cancer care. NPJ Precis Oncol, 2017. 1(1): p. 22.

7. Stang, A., et al., Diagnostic agreement in the histopathological evaluation of lung cancer tissue in a population-based case-control study. Lung Cancer, 2006. 52(1): p. 29–36.

8. Yu, K.H., et al., Predicting non-small cell lung cancer prognosis by fully automated microscopic pathology image features. Nat Commun, 2016. 7: p. 12474.

9. Komura, D. and S. Ishikawa, Machine Learning Methods for Histopathological Image Analysis. Comput Struct Biotechnol J, 2018. 16: p. 34–42.

10. Azuaje, F., Artificial intelligence for precision oncology: beyond patient stratification. NPJ Precis Oncol, 2019. 3: p. 6.

11. Rawat, W. and Z. Wang, Deep Convolutional Neural Networks for Image Classification: A Comprehensive Review. Neural Comput, 2017. 29(9): p. 2352–2449.

12. LeCun, Y., Y. Bengio, and G. Hinton, Deep learning. Nature, 2015. 521(7553): p. 436–44.

13. Litjens, G., et al., Deep learning as a tool for increased accuracy and efficiency of histopathological diagnosis. Sci Rep, 2016. 6: p. 26286.

14. Niazi, M.K.K., A.V. Parwani, and M.N. Gurcan, Digital pathology and artificial intelligence. Lancet Oncol, 2019. 20(5): p. e253–e261.

15. Kather, J.N., et al., Predicting survival from colorectal cancer histology slides using deep learning: A retrospective multicenter study. PLoS Med, 2019. 16(1): p. e1002730.

16. Tizhoosh, H.R. and L. Pantanowitz. Artificial Intelligence and Digital Pathology: Challenges and Opportunities, J Pathol Inform, 2018. 9: p. 38.

17. Liu, J., et al., An Integrated TCGA Pan-Cancer Clinical Data Resource to Drive High-Quality Survival Outcome Analytics. Cell, 2018. 173(2): p. 400–416.e11.

18. Institute, U.S.N.C. The Cancer Genome Atlas Program. [cited 2019; Available from: http://www.cancergenome.nih.gov/.

19. Institute, U.S.N.C. The Clinical Proteomic Tumor Analysis Consortium. Available from: https://proteomics.cancer.gov/.

20. Cooper, L.A., et al., PanCancer insights from The Cancer Genome Atlas: the pathologist’s perspective. J Pathol, 2018. 244(5): p. 512–524.

21. Mobadersany, P., et al., Predicting cancer outcomes from histology and genomics using convolutional networks. Proc Natl Acad Sci U S A, 2018. 115(13): p. E2970–e2979.

22. Yu, K.H., et al., Association of Omics Features with Histopathology Patterns in Lung Adenocarcinoma. Cell Syst, 2017. 5(6): p. 620–627.e3.

23. Zhong, T., M. Wu, and S. Ma, Examination of Independent Prognostic Power of Gene Expressions and Histopathological Imaging Features in Cancer. Cancers (Basel), 2019. 11(3).

24. Coudray, N., et al., Classification and mutation prediction from non-small cell lung cancer histopathology images using deep learning. Nat Med, 2018. 24(10): p. 1559–1567.

25. Srivastava, A., et al., Building trans-omics evidence: using imaging and ‘omics’ to characterize cancer profiles. Pac Symp Biocomput, 2018. 23: p. 377–387.

26. (CPTAC), N.C.I.C.P.T.A.C., National Cancer Institute Clinical Proteomic Tumor Analysis Consortium (CPTAC) collection proteomics and histology imaging datasets. 2018.

27. Simonyan, K. and A. Zisserman, Very Deep Convolutional Networks for Large-Scale Image Recognition. arXiv e-prints, 2014.

28. Subramanian, A., et al., Gene set enrichment analysis: A knowledge-based approach for interpreting genome-wide expression profiles. 2005. 102(43): p. 15545–15550.

29. NCI. GDC Data Portal - CPTAC-3. 2019; Available from: https://portal.gdc.cancer.gov/projects/CPTAC-3.

